# Differential effects of prediction error and adaptation along the auditory cortical hierarchy during deviance processing

**DOI:** 10.1101/2020.10.19.339234

**Authors:** Insa Schlossmacher, Jacky Dilly, Ina Protmann, David Hofmann, Torge Dellert, Marie-Luise Roth-Paysen, Robert Moeck, Maximilian Bruchmann, Thomas Straube

## Abstract

Neural mismatch responses have been proposed to rely on different mechanisms, including prediction error-related activity and adaptation to frequent stimuli. However, the cortical hierarchical structure of these mechanisms is unknown. To investigate this question, we used functional magnetic resonance imaging (fMRI) and an auditory oddball design with a suited control condition that enabled us to delineate the contributions of prediction error- or adaptation-related brain activation during deviance processing. We found that while prediction-error related processes increased with the hierarchical position of the brain area, adaptation declined. This suggests that the relative contribution of different mechanisms in deviance processing varies across the cortical hierarchy.

## Introduction

Detecting changes in our environment is a cornerstone of perception. A vast amount of research effort has been put into the investigation of neural correlates of deviance processing including neuroimaging (Kim, 2014) and electrophysiological approaches (Näätänen et al., 2011; Polich, 2007; Stefanics et al., 2014), showing increased activity to deviant stimuli in different variants of oddball designs. Recently, hierarchical predictive processing has been put forward as a theoretical underpinning for these deviance-related effects (Clark, 2013; Garrido et al., 2009; Stefanics et al., 2014; Winkler and Czigler, 2012). From this point of view, the increase of neural activation for unexpected rare stimuli compared to expected frequent ones stems from a process where a prediction is compared with the actual sensory input. If prediction and input do not match, as is the case when a rare – and thus unexpected – deviant stimulus is presented, a prediction error signal would be elicited. This prediction error signal would then be propagated upwards in the hierarchy and compared with the predictions of the next higher level and so forth, enabling efficient information processing (Clark, 2013; Stefanics et al., 2014).

However, while predictive processing offers a compelling explanation for deviance-related effects, it is not the only process that could be responsible for an observed difference between rare and frequent stimuli. Stimulus-specific adaptation (SSA) has also been proposed to play an important role during deviance processing (Jääskeläinen et al., 2004; May and Tiitinen, 2010; Nelken, 2014). In this theoretical framework, the difference between rare and frequent stimuli stems from habituation of neuronal responsiveness to the frequent stimulus, which thus elicits a smaller response compared to the non-adapted cells (fresh afferents) activated by the rare stimulus. In other words, deviance responses in typical oddball designs are not driven by genuine mismatch responses but by altered, i.e., reduced responses to the standard stimulus. While most work on SSA relies on electrophysiological studies in anmials, in humans, repetition suppression (RS) or fMRI adaptation (Barron et al., 2016; Larsson et al., 2016) represents a similar paradigm where neural activity after a stimulus repetition is reduced (Auksztulewicz and Friston, 2016; Barron et al., 2016). Similar to deviance responses in oddball paradigms, mechanisms relying on neuronal fatigue and predictive coding have been suggested to underlie repetition suppression (Auksztulewicz and Friston, 2016; Barron et al., 2016).

At first glance, the prediction error and the adaptation account seem opposing. However, there is evidence that both processes can be present in the brain to variable degrees in the same region or at the same time (Ishishita et al., 2019; Laufer et al., 2008; Opitz et al., 2005; Parras et al., 2017). One way to delineate the contribution of these two mechanisms to deviance effects is to compare the responses of deviant and standard stimuli to a control stimulus (Maess et al., 2007; Parras et al., 2017; Schröger and Wolff, 1996). The control stimulus is usually physically identical to the deviant but presented in a multi-standard paradigm where all stimuli are displayed with the same stimulus probability as the deviant. Comparing the deviant with the control thus offers a way to distill prediction error activity without being contaminated by adaptation. Additionally, comparing the control with the standard stimulus shows adaptation-related activity.

Using this experimental procedure, there is initial evidence from an electrophysiological study in rodents (Parras et al., 2017) that the relative contribution of prediction error vs. adaptation increases from subcortical areas to auditory cortex. In line with these results, two neuroimaging studies specifically targeting the auditory cortex showed the presence of both mechanisms within Heschl’s gyrus and superior temporal gyrus (STG) with an anterior-posterior gradient in humans (Laufer et al., 2008; Opitz et al., 2005). Furthermore, an electrocorticography (ECoG) study found that temporal areas were related to predictable changes, while frontal areas indexed unpredictable changes (Dürschmid et al., 2016). However, to date, there are no brain imaging studies that investigated the effects of prediction error vs. adaptation across all typical cortical brain areas commonly showcasing deviance-related effects (Kiehl et al., 2005; Kim, 2014). Besides primary and secondary auditory cortex, encompassing Heschl’s gyrus and STG, several frontal and parietal areas included in ventral and dorsal attention networks, as well as subcortical areas, reliably show increased activation to deviant vs. standard stimuli (Kiehl et al., 2005; Kim, 2014). Thus, deviance responses are processed on different hierarchical levels traversed during auditory processing from the thalamus to Heschl’s gyrus and STG and onwards to higher order association cortices (Li et al., 2019; Parras et al., 2017).

This hierarchical dimension of deviance processing, i.e., whether and how mechanisms vary depending on the cortical region, has not been investigated yet. In order to address this gap, we conducted a high-powered fMRI study (N = 54) using an oddball design with a suited control condition that allowed to delineate prediction error- and adaptation-related mechanisms in typical areas involved in auditory deviance processing in humans. Following prior research (Dürschmid et al., 2016; Parras et al., 2017), we hypothesized that on lower levels of the cortical hierarchy, adaptation-related activity will be more prominent, while on higher levels, activity should be driven by prediction errors.

## Methods

### Participants

Fifty-nine right-handed participants with normal hearing and no history of neurological or psychiatric illness took part in the experiment and were compensated with €10/h. Four participants had to be excluded due to excessive head movements (> 3 mm) during recording and one participant because of anatomical MRI anomalies. The remaining 54 participants (39 female) were aged from 18 to 33 (M = 23.20, SD = 3.04). The local ethics committee has approved the study and all procedures were carried out in accordance with the Helsinki declaration.

### Experimental procedure and stimulus material

Stimuli consisted of pure sine-tones of 600, 800, 1000, 1200, and 1400 Hz. Stimulus duration amounted to 100 ms, including rise/fall times of 10 ms. The sound volume was chosen to be easily audible during functional sequences but not unpleasantly loud. In oddball blocks, the 800 and 1200 Hz tones served as deviant and standard counterbalanced across participants. The probability of deviant to standard stimuli was 20:80 and stimulus presentation was pseudorandomized so that no two deviants were presented consecutively. In control blocks, all five tone stimuli were presented randomly with a probability of 20% (see Figure 1A). Throughout, participants’ task was to respond to target stimuli of 300 ms duration, which were randomly interspersed in the sequence. In total, 40 targets were included, the frequency of which conformed to the stimulus probability of the current block, i.e., all tone frequencies could be potential targets. Interstimulus intervals ranged from 1.04 to 17.25 s (M = 3.01, SD = 2.06) and were derived using optseq2 (Greve, 2009). Four different optseq sequences were computed and randomly assigned to two runs per participant, creating 12 different sequence combinations. In total, two runs of 250 stimuli (ca. 13 min each) were acquired and separated by a short break. One run could start with either an oddball or control block, which seamlessly changed to control or oddball block in the middle of the run. Whether the participants started with the oddball or control block was counterbalanced across participants. The second run was always presented in reverse order, e.g., if the first run was oddball-control, the second run was control-oddball. Before starting the experiment, a short practice block of about 1 min was presented in order to accustom the participants to the task. Stimuli in this practice block were structured like the experimental block that followed, i.e., they also included brief oddball and control sequences. At all times, a white fixation cross was presented on a black screen, and participants were asked to fixate during the run. Stimulus presentation and response collection were controlled by the software Presentation (Version 21.1, Neurobehavioral Systems, Albany, CA).

**Figure 1.**
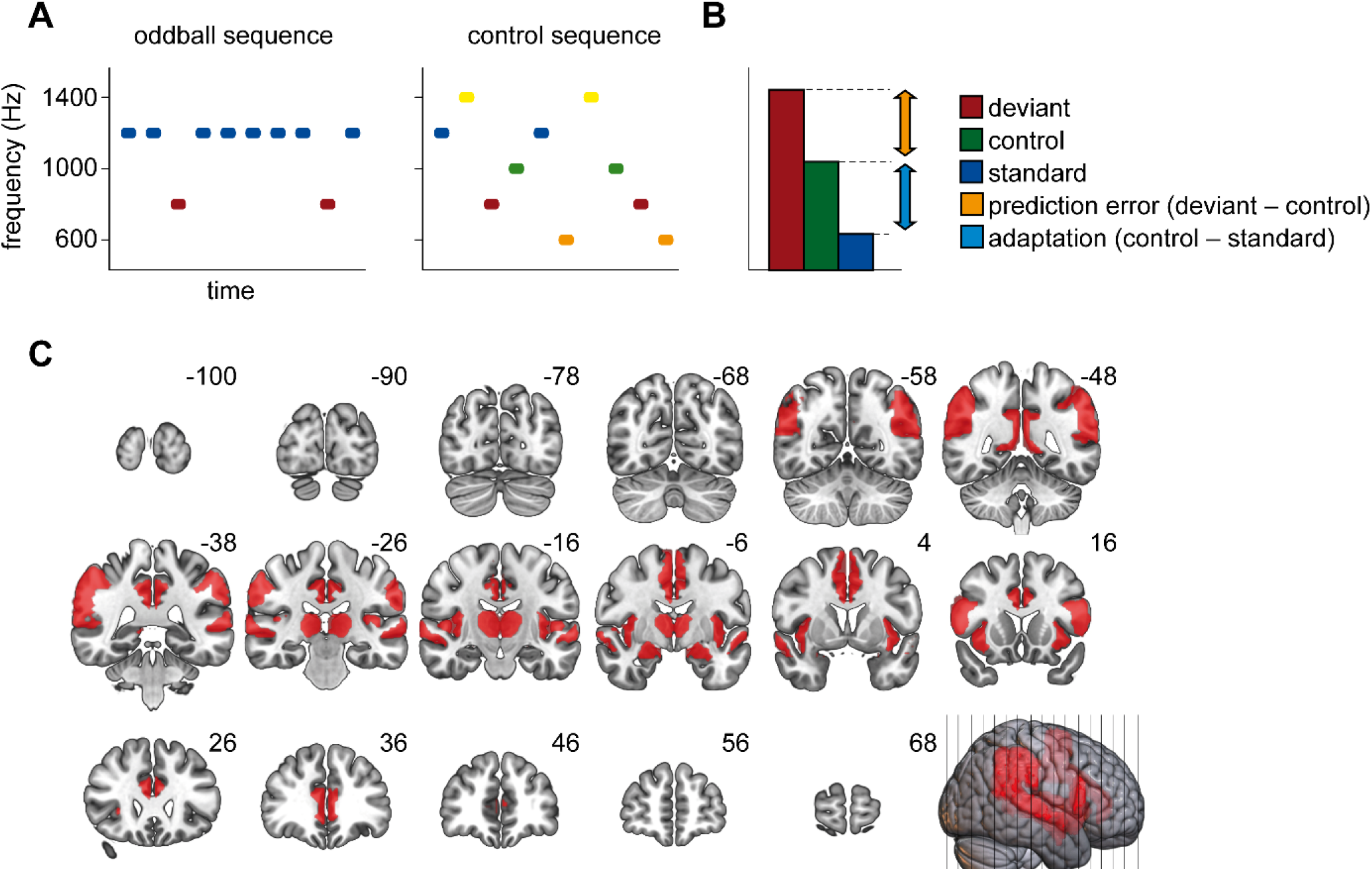
Experimental paradigm and mask for data analysis. (A) Schematic of oddball and control sequence. In the experiment, the frequencies of deviant and standard stimulus were counterbalanced across participants. (B) Decomposition of observed responses into prediction error-related and adaptation-related activity. (C) Illustration of the mask applied during the cluster-based permutation. Red areas were included in the mask.

### Data acquisition and preprocessing

A 3-Tesla Siemens Magnetom Prisma with a 20-channel Siemens Head Matrix Coil (Siemens Medical Systems, Erlangen, Germany) was used to aquire MRI data. In a first step, we obtained a high-resolution T1-weighted scan with 192 slices for anatomical localization and coregistration (repetition time (TR) = 2130 ms, echo time (TE) = 2.28 ms, flip angle (FA) = 8°, field of view (FOV) = 256 × 256 mm, voxel size = 1 × 1 × 1 mm). A shimming field was applied in order to minimize magnetic field inhomogeneity. Then, we recorded two functional datasets per participant (2 runs) consisting of 353 volumes and 42 slices each by means of a T2*-weighted echoplanar sequence sensitive to blood oxygenation level-dependent (BOLD) contrast (TR = 2300 ms, TE = 30 ms, FA = 90°, FOV = 216 × 216 mm, voxel size = 3 × 3 × 3 mm).

Preprocessing relied on SPM12 v7771 (Wellcome Department of Cognitive Neurology, London, UK) and the Data Processing & Analysis of Brain Imaging (DPABI) 4.3 toolbox (Yan et al., 2016) in MATLAB. We removed the first five data volumes to account for spin saturation effects. Then, slice-scan-time correction and realignment using a six-parameter (rigid body) linear transformation was performed. In a next step, we co-registered anatomical and functional images and segmented these into gray matter, white matter and cerebrospinal fluid. Finally, we normalized functional data to Montreal Neurological Institute (MNI) standard space using DARTEL (Ashburner, 2007), resampled it to 3 mm isotropic voxels and spatially smoothed it with an 8 mm full width at half maximum Gaussian kernel.

### Statistical Analysis

A general linear model (GLM) was estimated for each participant in the first-level analysis. In order to eliminate slow signal drifts, we used a high-pass filter with a cutoff of 128 seconds. We applied SPM’s pre-whitening method FAST (Corbin et al., 2018) to model autocorrelations as recommended by Olszowy and colleagues (2019). The GLM design matrix contained the onsets of deviants, standards, controls, targets and responses as predictors as well as six head movement parameters which represented predictors of no interest. We included two predictors accounting for the stimuli presented during the control condition. One used the onsets of the stimulus physically identical to the deviant (later compared with the deviant and standard), the other modeled the onsets of all other control stimuli. These onsets were then convolved with a 2-gamma hemodynamic response function to model the BOLD signal change for each predictor. Contrast images (deviant − standard) of the beta estimates were created for each participant for the second-level analysis.

In order to isolate mismatch-related activity in the second-level analysis, we used the cluster-based permutation as implemented in PALM (Winkler et al., 2014). The voxel-wise α amounted to .001; a cluster was deemed significant with α < .05. The number of permutations was set to 10000. Based on the recent meta-analysis of Kim (2014), the following areas of interest were identified and included in one mask based on the Harvard Oxford Atlas (Desikan et al., 2006): Heschl’s gyrus, superior temporal gyrus (STG), anterior and posterior cingulate cortex (ACC/PCC), supplementary motor area (SMA), inferior frontal junction (IFJ), inferior parietal lobule (IPL), temporo-parietal junction (TPJ), insula, thalamus and amygdala (see Figure 1C). Areas were chosen to correspond to the modality and task of the current study (auditory and task-irrelevant, see Kim, 2014 for details). In a second step, averaged betas of the found clusters were extracted for deviant, standard and control stimulus and z-standardized in order to account for differences in raw beta values between regions. For clusters comprising several distinct regions, we extracted the cluster averages using significant voxels encompassed in the original mask templates. Then, adaptation-related activity was computed as control – standard and prediction error-related activity as deviant – control (see Figure 1B). These differences were then subjected to a repeated-measures ANOVA to check whether adaptation-related and prediction error-related activity varied with brain region. When applicable, violations of sphericity were corrected using the Greenhouse-Geisser procedure and corrected *p*-values as well as 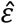–values are reported. Two-sided *t*-tests were used to follow up significant interactions. We additionally report Bayes Factors (BF), with BF_01_ denoting the evidence for the null hypothesis and BF_10_ the evidence for the alternative hypothesis. We use the conventions from Jeffreys (1961) to interpret the results of our Bayesian analyses.

### Data and code availability statement

Data and code are available on the Open Science Framework accessible via

https://osf.io/eufd6/.

## Results

### Behavioral Data

Performance on the duration task was very high, indexed by an average hit rate of 0.92 (SD = 0.15), an average false-alarm rate of 0.003 (SD = 0.007), and an average *d’* of 4.59 (SD = 0.77), indicating that participants were able to comply with the task easily. Furthermore, performance did not differ significantly between the oddball and control sequence (hit rate: *t*(53) = 0.66, *p* = .51, BF_01_ = 5.46; false alarm rate: *t*(53) = −0.81, *p* = .42, BF_01_ = 4.95; *d’*: *t*(53) = 0.15, *p* = .89, BF_01_ = 6.66).

### fMRI Data

#### Main effects of mismatch processing

The cluster-based permutation revealed bilateral clusters of significant mismatch processing in the auditory cortex, including STG and Heschl’s gyrus (right: *p* < .001; left: *p* < .001), the ACC/SMA (right: *p* = .01; left: *p* = .002), the IFJ (right: *p* = .003; left: *p* = .002) and the AI (right: *p* = .008; left: *p* = .006). Furthermore, a significant cluster was found in the left IPL (*p* = .009). Please see Figure 2 for a visualization of clusters and beta-values and Table 1 for peak coordinates, *t*-statistics, and number of voxels (k) of the effects.

**Figure 2.**
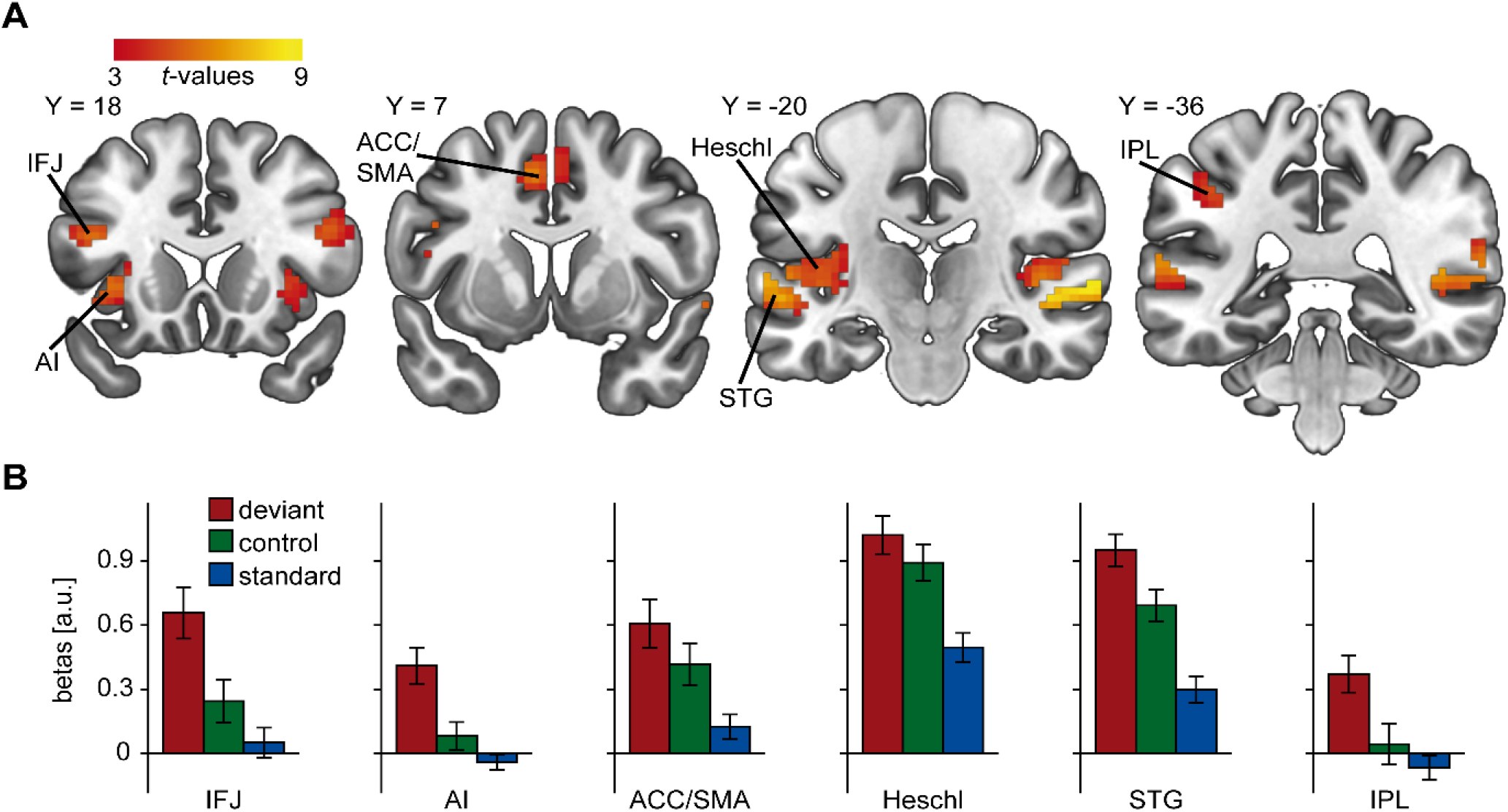
Auditory mismatch responses in the brain. (A) Clusters found in the cluster-based permutation comparing deviant and standard stimulus. (B) Mean beta values extracted from the clusters found. The deviant, standard and control stimulus are displayed. ACC/SMA: anterior cingulate cortex/supplementary motor area, AI: anterior insula, IFJ: inferior frontal junction, IPL: inferior parietal lobule, STG: superior temporal gyrus.

**Table 1.**
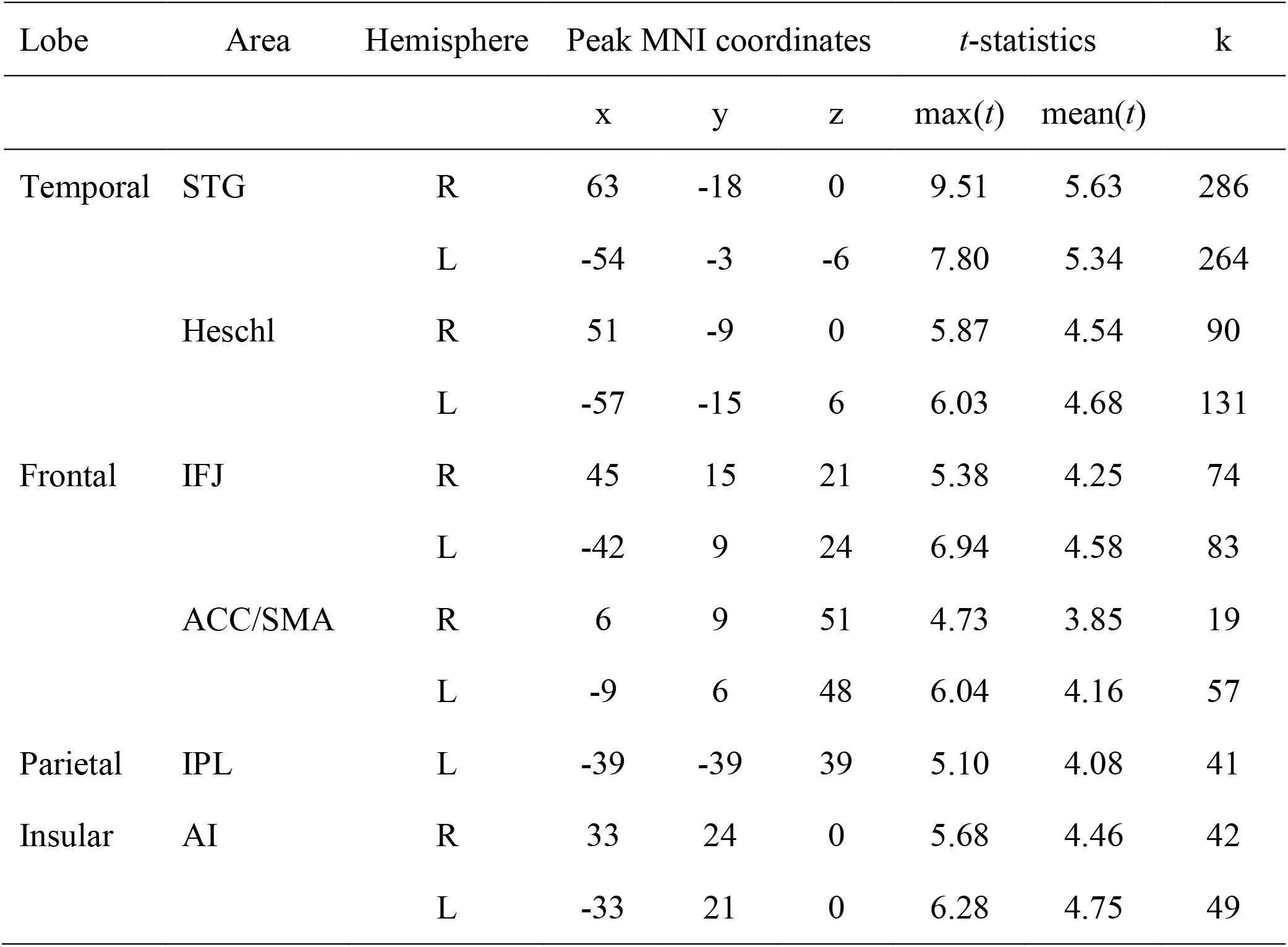
fMRI results of the oddball contrast. ACC/SMA: anterior cingulate cortex/supplementary motor area, AI: anterior insula, IFJ: inferior frontal junction, IPL: inferior parietal lobule, STG: superior temporal gyrus.

| Lobe | Area | Hemisphere | Peak MNI coordinates | | | $t$-statistics | | k |
| --- | --- | --- | --- | --- | --- | --- | --- | --- |
| | | | x | y | z | max($t$) | mean($t$) | |
| Temporal | STG | R | 63 | -18 | 0 | 9.51 | 5.63 | 286 |
|  |  | L | -54 | -3 | -6 | 7.80 | 5.34 | 264 |
|  | Heschl | R | 51 | -9 | 0 | 5.87 | 4.54 | 90 |
|  |  | L | -57 | -15 | 6 | 6.03 | 4.68 | 131 |
| Frontal | IFJ | R | 45 | 15 | 21 | 5.38 | 4.25 | 74 |
|  |  | L | -42 | 9 | 24 | 6.94 | 4.58 | 83 |
|  | ACC/SMA | R | 6 | 9 | 51 | 4.73 | 3.85 | 19 |
|  |  | L | -9 | 6 | 48 | 6.04 | 4.16 | 57 |
| Parietal | IPL | L | -39 | -39 | 39 | 5.10 | 4.08 | 41 |
| Insular | AI | R | 33 | 24 | 0 | 5.68 | 4.46 | 42 |
|  |  | L | -33 | 21 | 0 | 6.28 | 4.75 | 49 |

#### Mechanisms of mismatch processing

We found that the contribution of the different mechanisms varied depending on the brain region. The repeated-measures ANOVA indicated a significant main effect of area (*F*(5,265) = 2.54, *p* = .045, 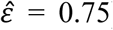) and a significant interaction of area and mechanism (*F*(5,265) = 3.28, *p* = .01, 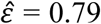), while the main effect of mechanism did not reach significance (*F*(1,53) = 0.03, *p* = .86). We found significant adaptation in Heschl’s gyrus (*t*(53) = 4.64, *p* < .001, BF_10_ = 847.85), STG (*t*(53) = 4.52, *p* < .001, BF_10_ = 588.07) and ACC/SMA (*t*(53) = 2.84, *p* = .006, BF_10_ = 5.41), while in the AI (*t*(53) = 1.81, *p* = .08, BF_01_ = 1.49), IPL (*t*(53) = 1.08, *p* = .29, BF_01_ = 3.90) and IFJ (*t*(53) = 1.57, *p* = .12, BF_01_ = 2.14), adaptation did not reach significance, see Figure 3A. In contrast, we observed prediction error in the STG (*t*(53) = 3.02, *p* = .004, BF_10_ = 8.36), AI (*t*(53) = 3.75, *p* < .001, BF_10_ = 57.97), IPL (*t*(53) = 2.93, *p* = .005, BF_10_ = 6.67) and IFJ (*t*(53) = 3.18, *p* = .002, BF_10_ = 12.42), while no significant prediction error was found in Heschl’s gyrus (*t*(53) = 1.39, *p* = .17, BF_01_ = 2.71) and ACC/SMA (*t*(53) = 1.42, *p* = .16, BF_01_ = 2.62).

**Figure 3.**
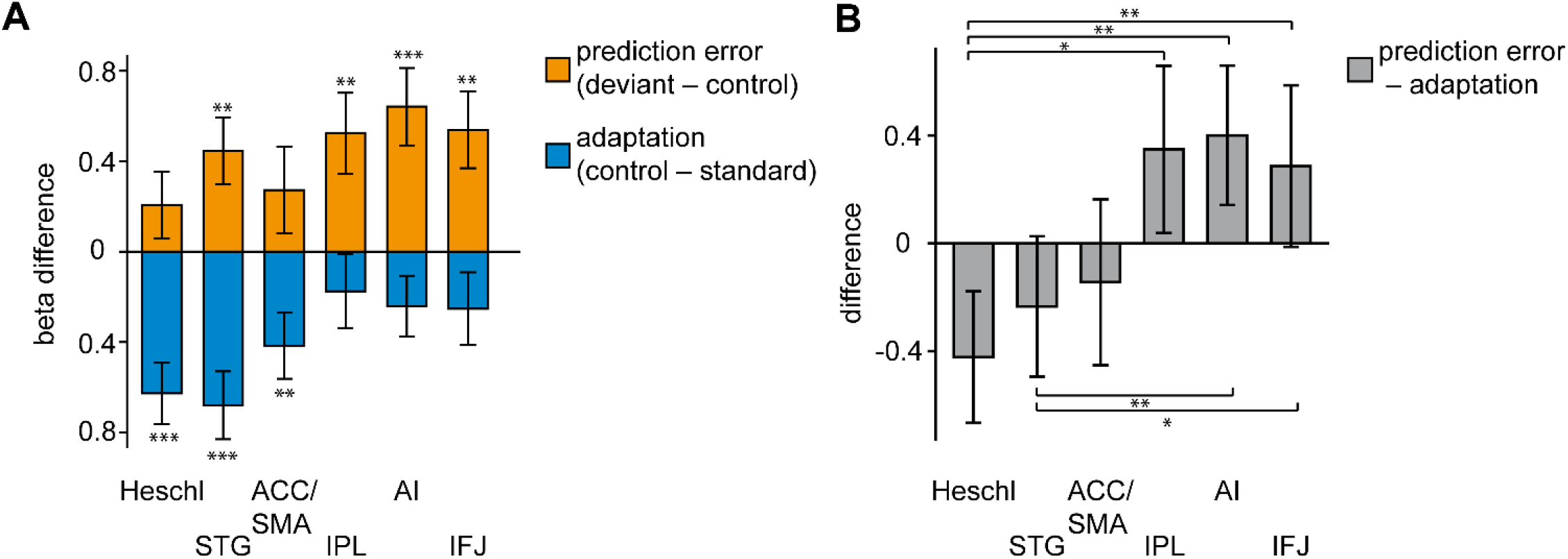
Mechanisms of mismatch generation. (A) Adaptation and prediction in different brain regions. Prediction-related activity is computed by substracting activity elicited by the control from the deviant. Adaptation-related activity is computed by substracting activity elicited by the standard from the control. Note that positive prediction error-related activity is plotted upwards and positive adaptation-related activity is plotted downwards. (B) Relative contribution of the mechanisms, negative values correspond to a surplus of adaptation in the corresponding brain area, while positive values correspond to a surplus of prediction error. Error bars depict standard errors of the mean. IFJ: inferior frontal junction, AI: anterior insula, ACC/SMA: anterior cingulate cortex/supplementary motor area, STG: superior temporal gyrus, IPL: inferior parietal lobule, Asteriks correspond to the *t*-tests reported in the main text with * p < .05, ** p < .01, *** p < .001.

In order to test how the relative contribution of the mechanisms differs between areas, we computed the difference between adaptation-related and prediction error-related activity. Mechanisms in Heschl’s gyrus differed significantly from AI (*t*(53) = −3.34, *p* = .002, BF_10_ = 18.75), IPL (*t*(53) = −2.48, *p* = .02, BF_10_ = 2.39) and IFJ (*t*(53) = −2.57, *p* = .01, BF_10_ = 2.89), STG differed significantly from AI (*t*(53) = −2.60, *p* = .01, BF_10_ = 3.13) and IFJ (*t*(53) = −2.38, *p* = .02, BF_10_ = 1.95). No other differences reached significance (all *p* > .05). See Figure 3B for a visualization of differences between cortical regions.

## Discussion

In this study, we investigated the contribution of different mechanisms to explain increased deviance-related brain activation in humans. We found a hierarchical increase of the relation of predictive mechanisms vs. adaptation from primary auditory cortex across secondary auditory cortex to higher frontal and parietal regions.

Our first analytical step, in which we compared oddball and standard stimuli, confirmed deviance-related activation in most of our ROIs. This includes bilateral Heschl’s gyrus and STG, which are commonly linked to auditory processing. Heschl’s gyrus encompasses the primary auditory cortex (Costa et al., 2011), while the STG is involved in higher-order auditory processing (Binder et al., 2000). The effects in both areas have been linked to the modulatory influences of the dorsal attention network (Kastner and Ungerleider, 2000), as well as preattentive change detection (Näätänen et al., 2007). Besides effects in sensory processing areas, we found several distinct clusters in frontal and parietal areas as well as the insula. In line with the meta-analytic results of Kim (2014) for task-irrelevant oddball studies, activations were mainly detected in regions belonging to the ventral attention network, which is involved in orienting attention to salient events and thus alerting the organism to environmental changes (Corbetta and Shulman, 2002; Sestieri et al., 2012). This includes the AI and ACC/SMA (Eckert et al., 2009; Yeo et al., 2011), which are also strongly involved in detecting salient events and initiating task-based attention (Goulden et al., 2014; Menon and Uddin, 2010; Shackman et al., 2011). Furthermore, we found strong bilateral IFJ activity. The IFJ is involved in the dorsal fronto-parietal network modulating goal-directed attention in sensory areas in a top-down fashion (Kim, 2014; Yeo et al., 2011), but also in a network activated by unexpected salient environmental changes (Sestieri et al., 2012). This duality fits with the suggestion that the IFJ presents a dynamic region integrating information from both dorsal and ventral networks (Asplund et al., 2010). In addition to these activations, we also found deviance-related effects in the left IPL, which is part of a fronto-parietal control network (Cole et al., 2013; Corbetta and Shulman, 2002; Yeo et al., 2011), probably involved in indexing expectancy violations (O’Connor et al., 2010).

These deviance-related effects could be explained to varying degrees by adaptation- and prediction error-related activity. We found significant contributions of adaptation in Heschl’s gyrus in line with previous research indicating sensory refractoriness effects in this area (Opitz et al., 2005), while contributions of prediction error did not reach significance. However, this does not mean that the primary auditory cortex does not show prediction error-related activation (Opitz et al., 2005), but that adaptation is – averaged across the deviance-related cluster – the primary driver of information processing. This point is further highlighted by the fact that we did not observe substantial evidence for the absence of an prediction error effect (BF_01_ < 3). In contrast to primary auditory cortex, prediction error-related activity was significantly observed in the STG in accordance with previous studies showing an increase of predictive processes along the auditory processing hierarchy (Laufer et al., 2008; Opitz et al., 2005; Parras et al., 2017).

Furthermore, in the AI, IFJ and IPL, a significant prediction error effect was found, but no significant adaptation effect. However, BFs in favor of the null hypothesis indicated substantial evidence only in the IPL, while in the AI and the IFJ, anecdotal evidence for the null was observed. This suggests that while prediction error is the primary driver of activity, adaptation effects cannot be ruled out with confidence. Nonetheless, the finding of prediction error as the primary mechanism fits well with studies showing a dominant role of these regions during various predictive processes (Allen et al., 2016; Dürschmid et al., 2016; Geuter et al., 2017; O’Connor et al., 2010; Siman-Tov et al., 2019). Surprisingly, we found no significant prediction error-related activity in ACC/SMA, probably due to the task-irrelevant oddball paradigm chosen here, which considerably differs from the paradigms linking ACC to predictive processing (Alexander and Brown, 2019). Furthermore, even though we used a large sample of participants, a further increase of sample size could probably alter results for ACC/SMA.

Comparing the relative contributions of prediction error and adaptation, we observed differences between areas mainly concerned with auditory processing and higher-order areas like the AI and IFJ. Thus, while prediction error is generated in auditory areas like the STG, its contribution to deviance-related effects increases on higher levels. These results are in line with the results of Dürschmid and colleagues (2016), who found an increase of predictive processing from temporal to frontal areas. Furthermore, this supposed hierarchical organization fits well with the results of Li and colleagues (2019). These authors reconstructed the temporal evolution of deviance-related responses by combining EEG and fMRI and found the information to flow from auditory cortex via the insula to the inferior frontal cortex. These findings are also in line with the proposal of hierarchical prediction error processing as a basic principle in the human brain (Clark, 2013; Heilbron and Chait, 2018). In this framework, auditory cortices are said to form a generative model of the acoustical environment, which is informed by predictions from higher level areas (Heilbron and Chait, 2018). We did not observe equal amounts of prediction error in all regions involved in deviance processing. At first glance, this seems to disagree with predictive coding theories that hypothesize the occurrence of error and prediction units on all cortical levels (Clark, 2013; Heilbron and Chait, 2018). However, if adaptation is considered as a means of the predictive brain to increase its precision, these findings can be integrated under the predictive coding framework (Garrido et al., 2009). Thus, in accordance with the results of Parras and colleagues (2017), we found evidence for a hierarchical organization of prediction errors during deviance processing.

While our study has many strengths, there are also limitations. In studies using electrophysiological measurements, SSA seems to be observed up to ISIs of 2 s (Ulanovsky et al., 2003). This is substantially shorter than the ISIs we chose here (M = 3 s). However, electrophysiological and hemodynamic evidence shows that in humans, deviance responses can still be observed using long ISIs (Bottcher-Gandor and Ullsperger, 1992; Juckel et al., 2012). Furthermore, even in the animal model, adaptation phenomena can be observed for ISIs up to 60 s in some cortical regions (Netser et al., 2011). Besides, while it is undisputed that the ISI is an important factor influencing deviance responses (Netser et al., 2011; Ulanovsky et al., 2003; Yarden and Nelken, 2017), the observation of differences between deviant and standard stimuli in a large number of different cortical regions in the current study speaks against the notion that deviance processing is completely abolished given ISIs > 2 s. Taken together, the current study shows that an oddball sequence can be implemented in an event-related fMRI paradigm with comparably long ISIs. Future studies should try to find ways to implement shorter ISIs while still maintaining the advantages of event-related fMRI.

As a second limitation, the equiprobable control might not have been the ideal control condition to delineate adaptation effects as controls were not as predictable as standards in the oddball sequence. Another control condition like the cascade control might have solved this issue by providing a predictable control stimulus (Ruhnau et al., 2012). However, first, while different results might be expected from equiprobable and cascade control in theory, empirical evidence suggests that both are equally suited to investigate mechanisms of change detection (Parras et al., 2017; Wiens et al., 2019). Second, we were interested in how the contributions of several mechanisms might change across the auditory processing hierarchy. Importantly, even if the control condition was not optimal, changes between different cortices should still be meaningful. Third, while treated as equally predictable, the standard oddball and the cascade control are not equal, as the next stimulus in the cascade control can be predicted with a probability of 100% while the standard in the oddball sequence cannot. Thus, while it is definitely desirable to use a cascade control in future studies, we think that in the current study, the use of the equiprobable control still allows us to draw conclusions on different mechanisms of deviance processing.

A third limitation is that we only investigated mechanisms of deviance processing during a condition where oddball stimuli had to be attended to but were not targets. Mechanisms predominantly observed during deviance processing might be modulated by task settings, thus experimentally combining task manipulations with fMRI might help better understand which brain regions vary in their predominant mechanisms and which do not. Besides, studies on repetition suppression have shown that the interplay between stimulus expectations and repetitions is complex, indicating that the influence of expectations on RS is not stable but changes with specific stimuli and conditions (Feuerriegel et al., 2018; Kovács et al., 2013; Tang et al., 2018; Todorovic et al., 2011; Utzerath et al., 2017). Thus, including different manipulations, e.g., of familiarity or higher-level expectations could further help in providing a comprehensive picture of hierarchical predictive processing in the brain. Furthermore, we only investigated the auditory modality. Future studies should also include other sensory stimulations. Finally, prospective research would profit from other experimental and analytical approaches for the investigation of different mechanisms of deviance processing.

## Conclusion

We observed deviance-related effects in a widespread network of different brain regions. The processes predominantly responsible for these effects varied depending on the hierarchical level of the brain region. We detected an increase in prediction error-related activity and a concurrent decrease of adaptation-related activity from lower to higher hierarchical areas. These results highlight hierarchical predictive processing in the human brain.

## Acknowledgments

The authors declare no competing financial interests.

